# *Maribellus comscasis* sp. nov., isolated from the deep-sea cold seep

**DOI:** 10.1101/2021.01.06.425552

**Authors:** Rikuan Zheng, Chaomin Sun

**Affiliations:** CAS Key Laboratory of Experimental Marine Biology & Center of Deep Sea Research, Institute of Oceanology, Chinese Academy of Sciences, Qingdao, China; Laboratory for Marine Biology and Biotechnology, Qingdao National Laboratory for Marine Science and Technology, Qingdao, China; College of Earth Science, University of Chinese Academy of Sciences, Beijing, China; Center of Ocean Mega-Science, Chinese Academy of Sciences, Qingdao, China

**Keywords:** *Maribellus comscasis*, 16S rRNA gene sequence, polysaccharides, deep-sea cold seep

## Abstract

A facultatively anaerobic, Gram-stain-negative, non-motile, curved rod-shaped bacterium, designated WC007^T^, was isolated from the deep-sea cold seep, P. R. China. Strain WC007^T^ was found to grow at temperatures from 28 to 37 °C (optimum, 30 °C), at pH values between pH 6.0 and 8.0 (optimum, pH 7.0) and in 0-5.0% (w/v) NaCl (optimum, 1.0%). The major fatty acids (>10.0%) were iso-C_15:0_, C_16:0_, summed feature 3 and summed feature 8. The major isoprenoid quinone was MK-7. Predominant polar lipids were phosphatidylethanolamine, one unidentified phospholipid, one unidentified aminolipid and one unidentified lipid. The G+C content of the genomic DNA was 38.38%. The average nucleotide identity (ANIb and ANIm), amino acid identity (AAI), the tetranucleotide signatures (Tetra) and *in silico* DNA-DNA hybridization (*is*DDH) similarities between the genome sequences of isolate WC007^T^ and *Maribellus luteus* XSD2^T^ were 70.11%, 84.94%, 71.0%, 0.92022 and 20.40%, respectively, indicating that strain WC007^T^ was distinguished from *M. luteus*. Phylogenetic analysis based on 16S rRNA gene sequences placed strain WC007^T^ within the genus *Maribellus* and showed the highest similarity to strain XSD2^T^ (95.70%). In combination of the results of phylogenetic analysis and phenotypic and chemotaxonomic data, strain WC007^T^ was considered to represent a novel species of the genus *Maribellus*, for which the name *Maribellus comscasis* sp. nov. is proposed. The type strain is WC007^T^ (=KCTC 25169^T^ = MCCC 1K04777^T^). The available of the genome sequence of strain WC007^T^ would be helpful in understanding the degradation mechanism of difficult-to-degrade polysaccharides.

---

The family Prolixibacteraceae was originally described and classified under the order Bacteroidales [1], class Bacteroidia, phylum Bacteroidetes, and the genus *Prolixibacter* is the type genus [2]. After that, the families Prolixibacteraceae, Marinilabiliaceae and Marinifilaceae were transferred to the order Marinilabiliales in 2016 [3]. At the time of writing, 11 genera including *Aquipluma*, *Meniscus*, *Puteibacter*, *Sunxiuqinia*, *Prolixibacter*, *Mangrovibacterium*, *Draconibacterium*, *Mariniphaga*, *Tangfeifania*, *Roseimarinus* and *Maribellus* had been identified within the family Prolixibacteraceae [4–6]. Members of the family Prolixibacteraceae were isolated from various habitats, such as coastal sediment [7, 8], crude oil [9], mangrove sediment [1], freshwater lake [5], and marine sediments [10]. The new genus *Maribellus* within the family Prolixibacteraceae was proposed recently [4], and *Maribellus luteus* XSD2^T^, isolated from coastal seawater, was the only strain in the genus *Maribellus*. The predominant respiratory quinone of strain XSD2^T^ was MK-7, which is the most frequently identified respiratory quinone in bacteria belonging to the family Prolixibacteraceae [4].

The phylum Bacteroidetes, specialized on polysaccharide degradation, was the most abundant group of bacteria in the ocean after Proteobacteria and Cyanobacteria [11]. Marine Bacteroidetes were commonly assumed to have a key role in degrading phytoplankton polysaccharides [12], which had a great number and diversity of carbohydrate-active enzymes (CAZymes) [13]. The CAZymes are categorized into families of glycoside hydrolases (GHs), glycoside transferases (GTs), carbohydrate-binding modules (CBMs), carbohydrate esterases (CEs), polysaccharide lyases (PLs), sulfatases (targeting sulfated polysaccharides) plus a range of auxiliary enzymes [12, 14, 15]. And the components of Bacteroidetes for the degradation of polysaccharides was often encoded in distinct polysaccharide utilization loci (PULs) [16], which were strictly regulated by the gene clusters that encode CAZymes and protein ensembles required for the degradation of complicated carbohydrates. Besides the various substrate-specific CAZymes genes, Bacteroidetes PULs also contain genes encoding SusCD-like proteins that are extracellular lipoproteins and integral membrane beta-barrels termed TonB-dependent transporters [17]. In addition, PULs of marine Bacteroidetes are frequently found many sulfatases [15, 18–21], because abundant polysaccharides from marine algae were often sulfated. So far most functional studies of Bacteroides polysaccharide degradation have been conducted in human gut Bacteroides [22], however, only a few studies have investigated polysaccharide degradation by marine Bacteroides, especially the deep-sea variety [23].

In this study, a facultatively anaerobic strain WC007^T^, belonging to the *Maribellus* genus, was isolated from deep-sea sediment at a depth of 1,146 m in the cold seep (22° 06’ 58.598’’ N 119° 17’ 07.322’’ E) as described previously [24], P. R. China. Strain WC007^T^ was isolated from an enrichment medium containing (per liter of seawater): 1.0 g NaHCO_3_, 1.0 g CH_3_COONa, 1.0 g NH_4_Cl, 0.5 g KH_2_PO_4_, 0.2 g MgSO_4_·7H_2_O, 1.0 g polysaccharide (cellulose, pectin and xylan), 1.0 mL 0.1 % (w/v) resazurin, 0.7 g cysteine hydrochloride (pH 7.0) at atmospheric pressure and the medium was prepared anaerobically as previously described [25]. The cultures were repeatedly purified by using the Hungate roll-tube method in the medium containing 1.5% (w/v) agar. After incubation for 7 days, several colonies were picked by sterilized bamboo skewers and then harvested and cultured in the liquid medium. The process of isolation was repeated several times until the isolates were deemed to be axenic. The purity of this isolate was confirmed routinely by transmission electron microscopy (TEM) and by repeated sequencing of the 16S rRNA gene. Then the single colony was transferred to a new medium (ORG) for further culture. The ORG medium contained (L^−1^): 1.0 g NaHCO_3_, 1.0 g NH_4_Cl, 1.0 g CH_3_COONa, 0.2 g MgSO_4_·7H_2_O, 0.5 g KH_2_PO_4_, 1.0 g yeast extract, 1.0 g peptone, 0.7 g cysteine hydrochloride, 1 mL 0.1% (w/v) resazurin; the pH was adjusted to 7.0 [26]. The isolate was cultured in ORG broth and maintained at −80 °C as a suspension in ORG supplemented with glycerol (20 %, v/v).

Genomic DNA was extracted from strain WC007^T^ cultured for 7 d at 30 °C using a bacteria genomic DNA kit (Takara, Japan). The whole genome sequencing (WGS) of strain WC007^T^ was performed on the nanopore sequencing technology platform with the Oxford Nanopore MinION (Oxford, UK) and Illumina MiSeq sequencing platform (San Diego, USA). The experimental process was implemented in accordance with the standard protocol provided by Oxford Nanopore Technologies (Oxford, UK). And the library building includes the following four steps: (1) High quality genomic DNA was extracted, and then Nanodrop, Qubit and 0.35 % agarose gel electrophoresis were used for purity, concentration and integrity inspection, respectively; (2) The large fragments of DNA were recovered by the BluePippin automatic nucleic acid recovery system; (3) Library construction was conducted by using the SQK-LSK109 connection Kit (Japan), which including DNA damage repair and terminal repair, magnetic bead purification, ligation of sequencing adapters and magnetic bead purification. (4) After the library construction, computer sequencing will be performed and Canu V1.5 software was used to assemble the filtered subreads [27]. Finally, Pilon software was used to correct the assembled genome with second-generation data to obtain the final genome with higher accuracy [28].

The whole genome of strain WC007^T^ had been deposited at GenBank under the accession number CP046401. The genome size of strain WC007^T^ was 7,811,310 bp with a DNA G+C content of 38.38%. The number of contigs was 1, the total of N50 was 7,811,310 and the sequencing depth was 50.0×. Annotation of the genome of strain WC007^T^ consisted of 6,176 coding sequences that included 54 RNA genes (6 rRNA genes, 45 tRNA genes and 3 other ncRNAs). And we checked the authenticity of the genome using the QUAST-5.0.2 software. The results showed that the genome assembly quality of WC007^T^ was high. In the genome of WC007^T^, genes encoding SusC, SusD, CAZymes and sulfatase were ubiquitously distributed, strongly indicating it possesses potentials for polysaccharide degradation. As measures of relatedness between strain WC007^T^ and closely related strains, the genome relatedness values were calculated by several approaches: Average Nucleotide Identity (ANI) based on the MUMMER ultra-rapid aligning tool (ANIm), ANI based on the BLASTN algorithm (ANIb), the tetranucleotide signatures (Tetra), and in silico DNA–DNA similarity. ANIm, ANIb, and Tetra frequencies were calculated, using JSpecies WS (http://jspecies.ribohost.com/jspeciesws/) [29]. The recommended species criterion cut-offs were used; 95% for the ANIb and ANIm and 0.99 for the Tetra signature [30]. The amino acid identity (AAI) values were calculated by AAI-profiler (http://ekhidna2.biocenter.helsinki.fi/AAI/) [31]. The *in silico* DNA-DNA similarity (*is*DDH) values were calculated by the Genome-to-Genome Distance Calculator (GGDC) (http://ggdc.dsmz.de/) [32]. The *is*DDH results were based on the recommended formula 2, which is independent of genome size and, thus, is robust when using whole-genome sequences.

From ANI calculations, it could be observed that the comparison between strain WC007^T^ and three closely relatives were noticeably lower (ANIb: 70.11-70.51% and ANIm: 83.71-84.94%) than the 95% cut-off proposed for bacterial species delineation. The AAI values of 71.0-71.7% between strain WC007^T^ and closely relatives were obtained, which were also far below the 95% cut-off value generally recommended for species differentiation. The Tetra values were also lower (0.92022-0.9496) than the 0.99 cut-off proposed for bacterial species differentiation. Finally, the *is*DDH values were significantly lower (18.40-20.40%) than the 70% cut-off value generally recommended for species differentiation. The related genome comparison data were listed in the Table S1.

Genomic DNA extraction and PCR amplification of the 16S rRNA gene of strain WC007^T^ were carried out as described by Hetharua et al [33]. The PCR product was cloned into the vector pMD-19T (TaKaRa, Japan), sequenced and then compared to the 16S rRNA gene sequence extracted from the genome. The results exhibited 99.9 % sequence similarity. The whole full-length 16S rRNA gene sequence (1,516 bp) of WC007^T^ was obtained from the genome, which had been deposited in the GenBank database (accession number MN096653). Moreover, the 16S rRNA gene sequences of related taxa were obtained from NCBI (www.ncbi.nlm.nih.gov/). A sequence similarity calculation using the NCBI server indicated the closest relatives of strain WC007^T^ were *Maribellus luteus* XSD2^T^ (95.70%), *Mariniphaga sediminis* SY21^T^ (93.44%) and *Draconibacterium sediminis* MCCC 1A00734^T^ (92.99%). Phylogenetic analysis was performed using the software MEGA version 6.0 [34]. Phylogenetic trees were constructed by the neighbor-joining algorithm [35], maximum Likelihood [36] and minimum-evolution methods [37]. The numbers above or below the branches are bootstrap values based on 1,000 replicates. Phylogenetic analyses based on the 16S rRNA and genome sequences showed strain WC007^T^ belonged to the genus *Maribellus* and formed an independent phyletic line with the type strain *Maribellus luteus* XSD2^T^ (Fig. 1 and Fig. S1). Therefore, we propose strain WC007^T^ to be classified as the type strain of a novel species in the genus *Maribellus*, for which the name *Maribellus comscasis* sp. nov. is proposed. On the basis of the phylogenetic results, *Maribellus luteus* XSD2^T^ (=MCCC 1H00347^T^) was selected as the closest recognized neighbor of strain WC007^T^ and was used as a reference strain in most of the subsequent phenotypic tests.

**Fig. 1.**
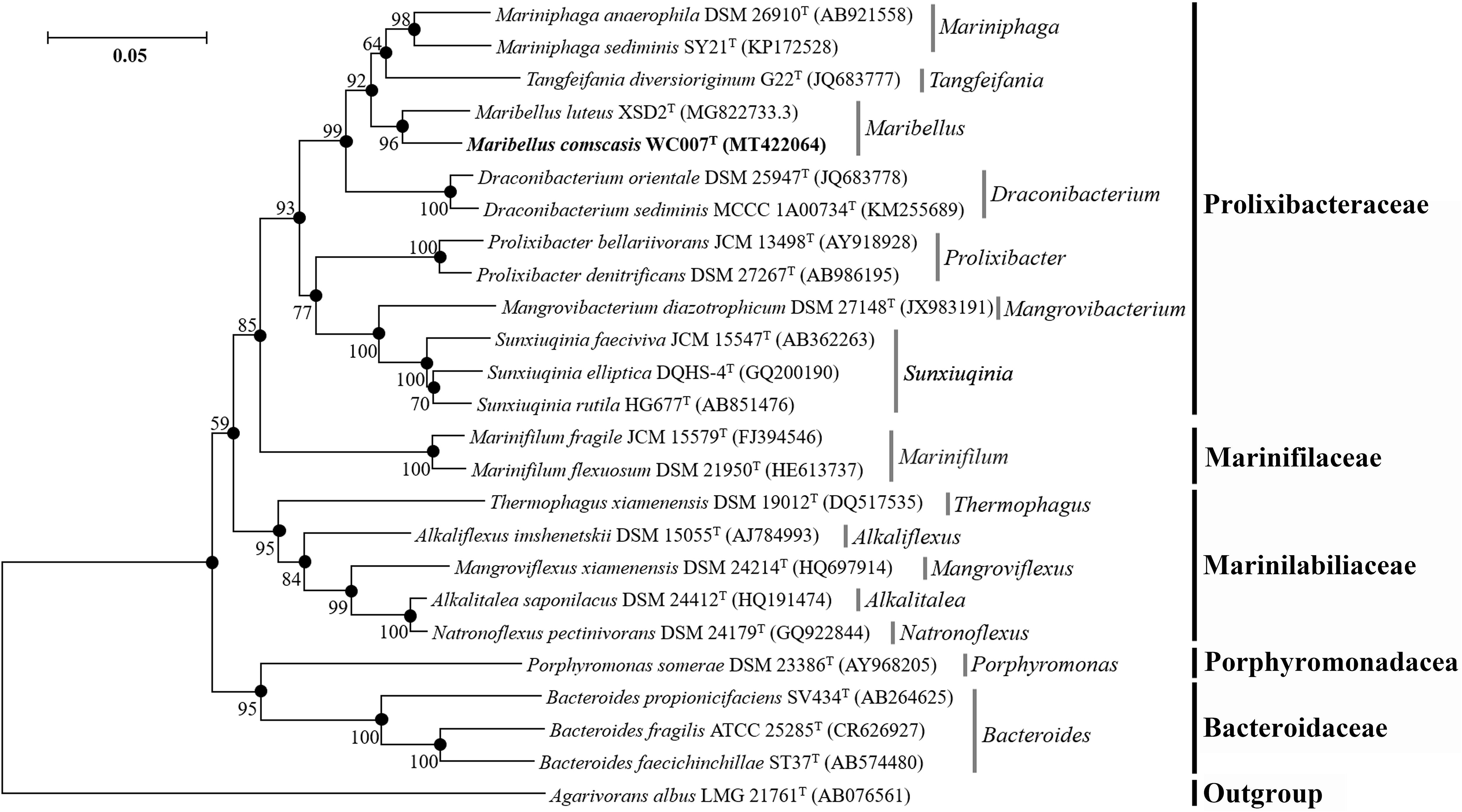
Neighbor-joining tree based on 16S rRNA gene sequences showing the position of the novel strain WC007^T^ among bacterial members of the family Prolixibacteraceae and other closely related families. The filled circles indicate branches of the tree that were also formed using the maximum-likelihood and minimum-evolution methods. Numbers at branching points are bootstrap values (expressed as percentages of 1000 replications). The access number of each 16S rRNA is indicated after the strain’s name. The sequence of *Agarivorans albus* LMG 21761^T^ is used as an outgroup. Bar, 0.05 substitutions per nucleotide position.

To observe the morphological characteristics of *M. comscasis* WC007^T^, cells were examined using transmission electronic microscopy (TEM) (HT7700; Hitachi, Japan) with a JEOL JEM 12000 EX (equipped with a field emission gun) at 100 kV. The cells suspension of *M. comscasis* WC007^T^ was washed with Milli-Q water and centrifuged at 5,000 *g* for 5 min. Subsequently, the sample was taken by immersing copper grids coated with a carbon film for 20 min in the bacterial suspensions and washed for 10 min in distilled water and dried for 20 min at room temperature [38]. Strain WC007^T^ was Gram-stain-negative and showed a curved rod-shaped, 1.0-3.5×0.5-0.8 μm in size, which had no flagellum as indicated by TEM (Fig. 2).

**Fig. 2.**
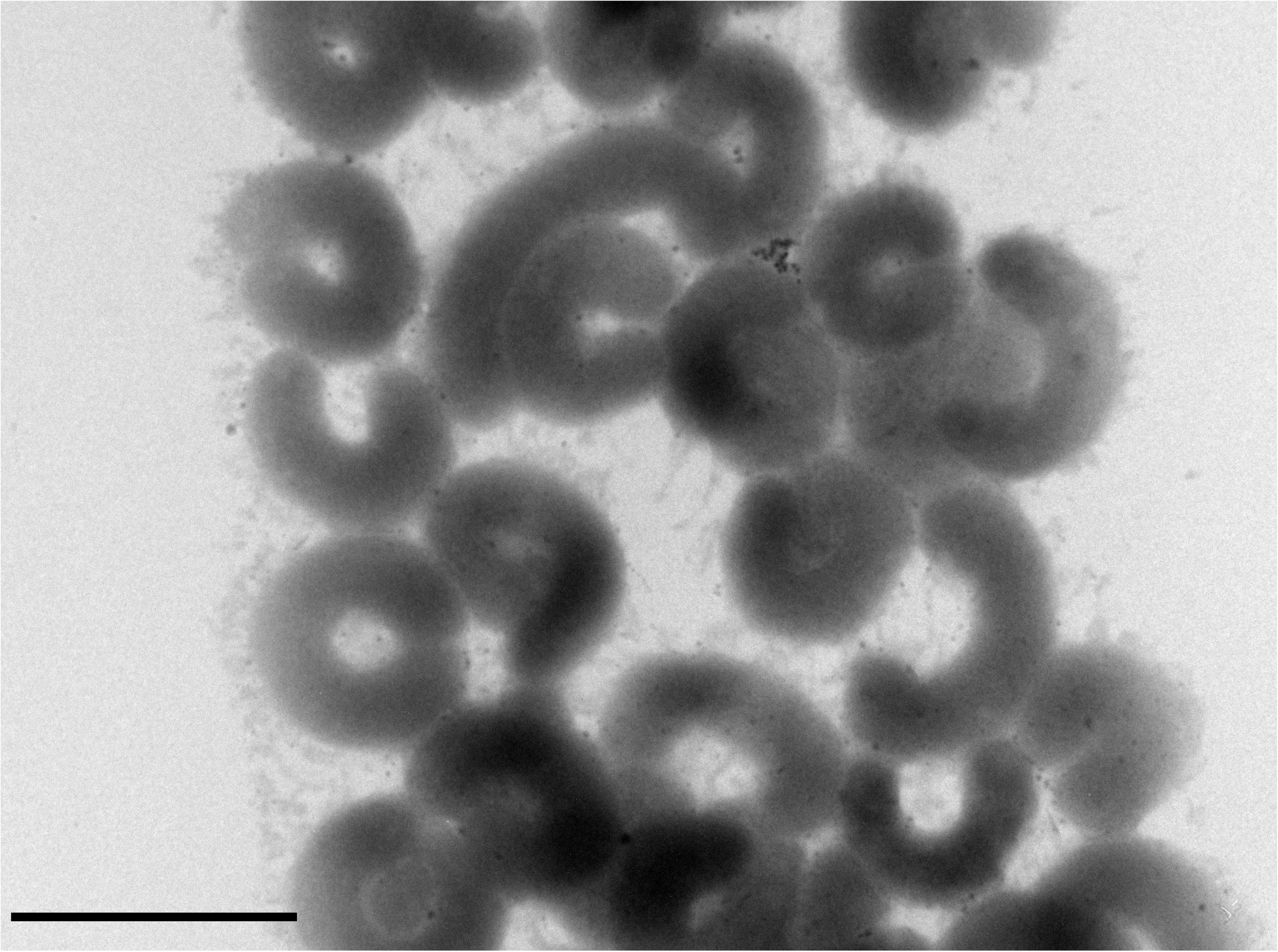
Transmission electron microscopy (TEM) observation of a negatively stained culture of strain WC007^T^. Bar is 2 μm.

For phenotypic characteristics comparison, the temperature, pH and NaCl concentration ranges for the growth of strain WC007^T^ were determined in duplicate experiments using the ORG medium as described above. Water baths and thermostatic incubators were used to incubate bacterial cultures from 4 to 80 °C. The pH of the culture medium was adjusted by 6 M HCl for low pH and 10% NaHCO_3_ (w/v) for high pH. The pH range for growth was tested from pH 3.0 to pH 11.0 (initial pH at 30 °C) with increments of 0.5 pH units. Salt tolerance was determined by directly weighing NaCl (0-100 g L^−1^) into the Hungate tubes before packaging the autotrophic medium. Catalase activity was determined by observation of bubble production after the application of 3% (v/v) hydrogen peroxide solution. Oxidase activity was evaluated by the oxidation of 1% (w/v) tetramethyl p-phenylenediamine. Substrate utilization was tested at atmospheric pressure in duplicate, using Hungate tubes containing basal medium contained (L^−1^): 1.0 g NaHCO_3_, 1.0 g NH_4_Cl, 1.0 g CH_3_COONa, 0.2 g MgSO_4_·7H_2_O, 0.5 g KH_2_PO_4_, 0.7 g cysteine hydrochloride, 1 mL 0.1% (w/v) resazurin. Single substrate (including cellulose, pectin, xylan, glucose, acetate, maltose, butyrate, fructose, glycine, ethanol, formate, lactate, sucrose, sorbitol, D-mannose) was added from sterile filtered stock solutions to the final concentration at 20 mM, respectively. Cell culture without adding any other substrates was used as a control. The pH was adjusted to 7.0 with NaOH. The cultures were incubated at 30 °C for 7 days and then determined by spectrophotometry at 600 nm. For each substrate was repeated three times. All tested substrates were listed in the species description.

The detailed physiological characteristics that differentiate the *Maribellus luteus* XSD2^T^ were listed as Table 1. Strain WC007^T^ required NaCl for growth and growth was observed at 0-5.0% NaCl (optimum: 1.0% NaCl). And growth occurred at 28-37 °C (optimum, 30 °C) and at pH 6.0-8.0 (optimum, pH 7.0). In addition, the positive activities of acetate, maltose, fructose, ethanol, formate, lactate, sorbitol and D-mannose in strain WC007^T^ distinguished those in strain XSD2^T^. The differential phenotypic characteristics between strain WC007^T^ and the closely related type strain XSD2^T^ are shown in the Table 1. Overall, the phenotypic characterization supports that strain WC007^T^ represents a novel species of the genus *Maribellus*.

**Table 1.**
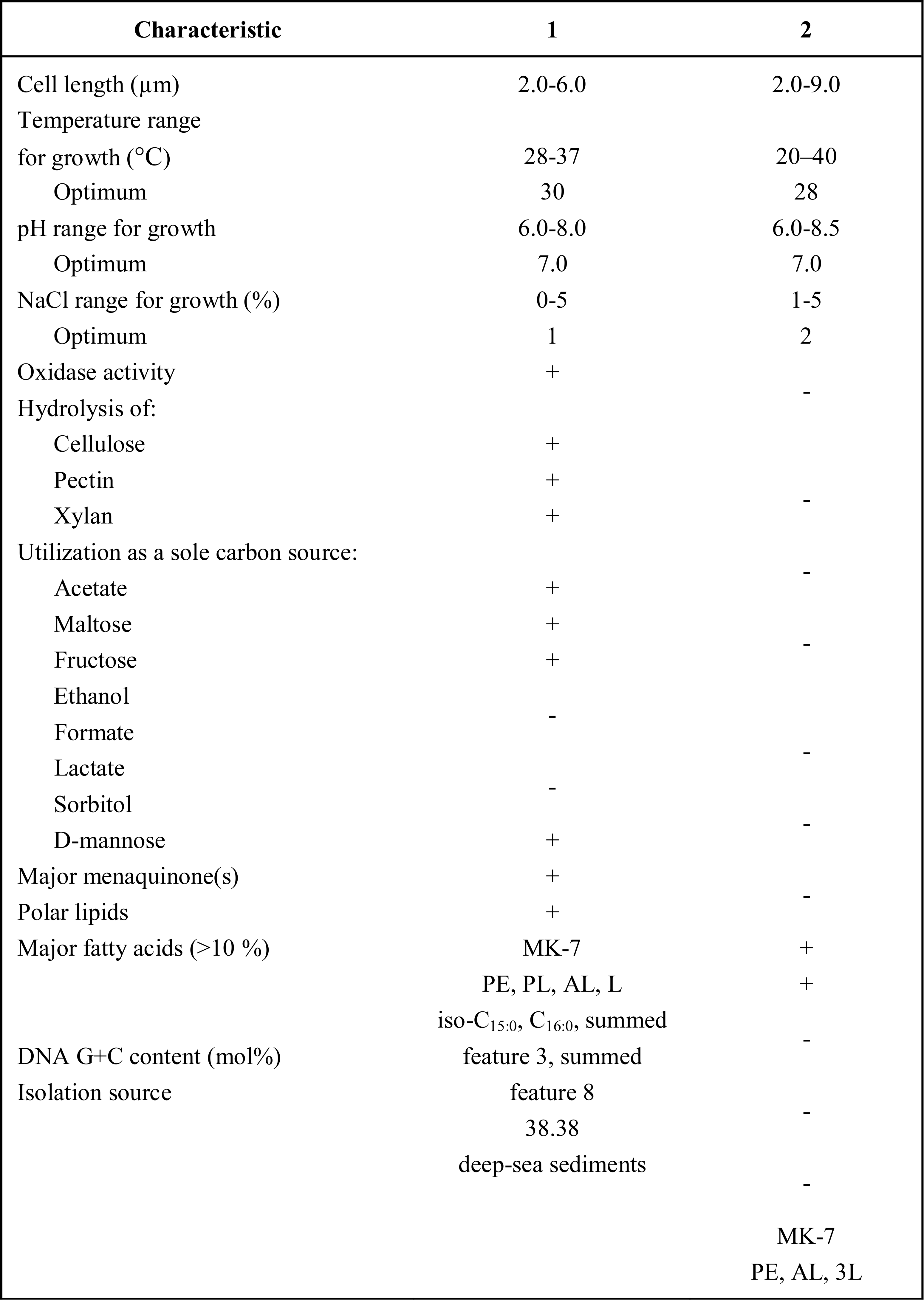
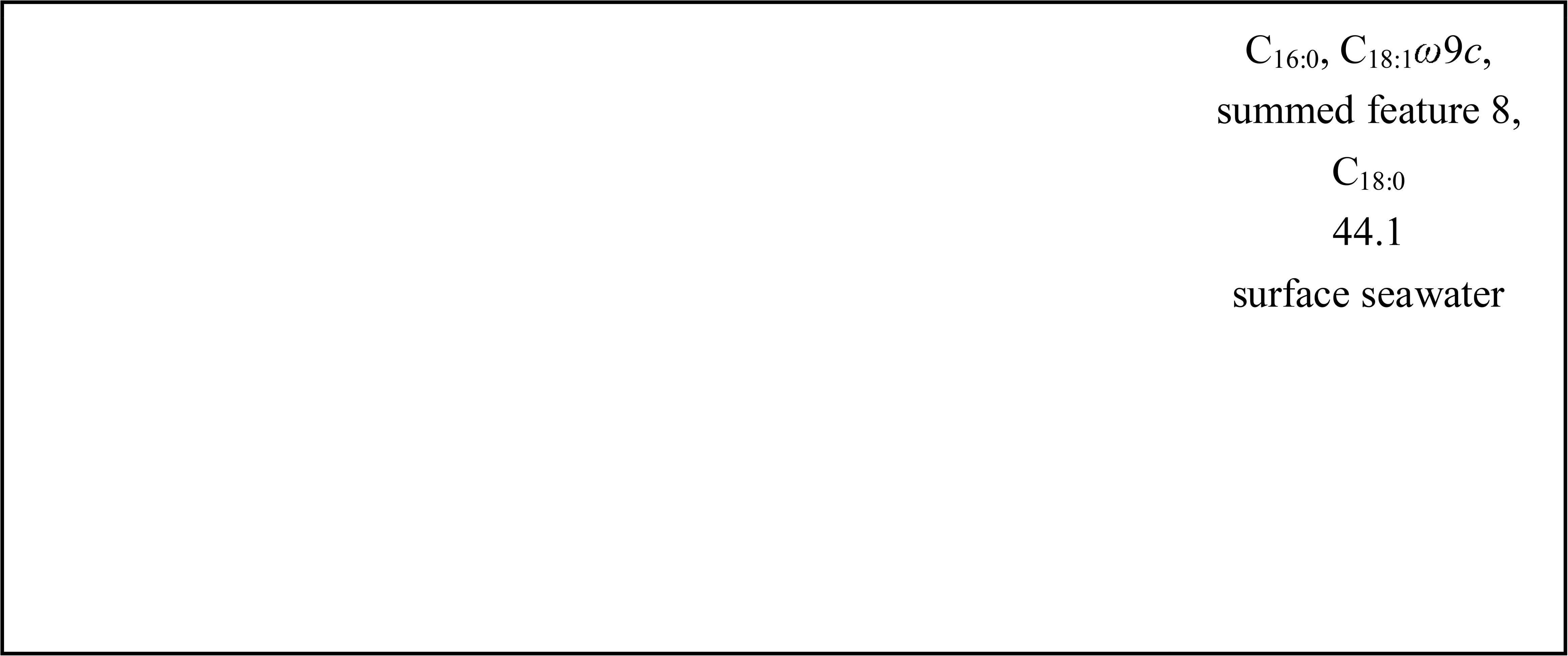
Differential physiological characteristics of the novel strain WC007^T^ and its closest related type strain *Maribellus luteus* XSD2^T^. Strains: 1, WC007^T^ (all data from this study); 2, *Maribellus luteus* XSD2^T^ (all data from this study except DNA G+C content and polar lipids). +, Positive result or growth; -, negative result or no growth. *Summed features are groups of two or three fatty acids that could not be separated by GLC using the MIDI system. Summed feature 8 contains C_18:1_*ɷ*7*c* and/or C_18:1_*ɷ*6*c*. Summed feature 8 contains C_16:1_*ɷ*7*c*/C_16:1_*ɷ*6*c* and/or C_16:1_*ɷ*6*c*/C_16:1_*ɷ*7*c*.

**Table 2.**
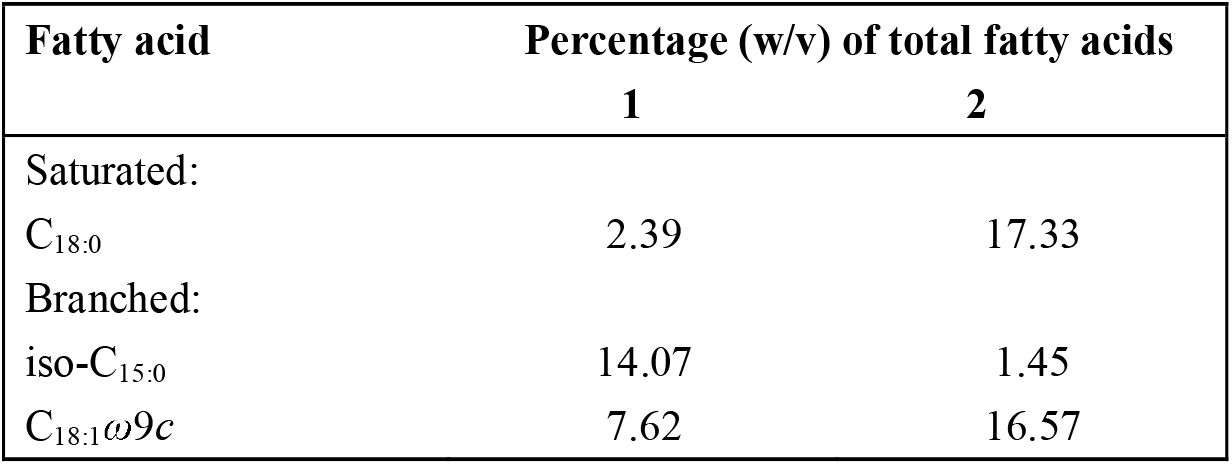
Percentages of fatty acids useful for distinguishing WC007^T^ from its closest relative *Maribellus luteus* XSD2^T^. Strains: 1, WC007^T^ (all data from this study); 2, *Maribellus luteus* XSD2^T^ (all data from this study).

For chemotaxonomic analysis, cells of WC007^T^ were cultured and collected under the same conditions unless stated otherwise, with the closely related type strains were grown on the ORG solid medium for 7 d at 30 °C under the same condition. Cells of WC007^T^ and the closely related type strain were harvested from cultures at the mid-exponential phase of growth and freeze-dried. Cellular fatty acids were extracted and determined from dried cells by using GC (model 7890A, Agilent, USA) according to the protocol of the Sherlock Microbial Identification System [39]. Polar lipids were extracted and determined as described by Tindall et al [40].

The predominant respiratory quinone of strain WC007^T^ was MK-7, which is the most frequently identified respiratory quinone in bacteria belonging to the family Prolixibacteraceae [4]. The major cellular fatty acids (>10.0 %) in strain WC007^T^ were iso-C_15:0_, C_16:0_, summed feature 3 and summed feature 8. The amount of iso-C_15:0_ in strain WC007^T^ (14.07%) was higher than that found in strain XSD2^T^ (1.45%), while the amount of C_18:0_ and C_18:1_ω9*c* in strain WC007^T^ (2.39%, 7.62%) was lower than that found in strain XSD2^T^ (17.33%, 16.57%), respectively. The major polar lipids in strain WC007^T^ were phosphatidylethanolamine, one unidentified phospholipid, one unidentified aminolipid and one unidentified lipid (Fig. S2). Strain WC007^T^ was apparently different from the type strain XSD2^T^ by the presence of one unidentified phospholipid.

In summary, phylogenetic analysis of strain WC007^T^ based on 16S rRNA gene sequence similarities confirmed the distinctness of strain WC007^T^ from the closely related strain XSD2^T^. Moreover, based on a polyphasic taxonomic approach and several phenotypic characteristics, strain WC007^T^ distinguished from *Maribellus luteus* XSD2^T^, the only recognized species of the genus *Maribellus*. Therefore, we propose that strain WC007^T^ was classified as the type strain of a novel species in the genus *Maribellus*, for which the name *Maribellus comscasis* sp. nov. is proposed.

## Description of *Maribellus comscasis* sp. nov.

*Maribellus comscasis* (com.sca’sis. L. gen. pl. n. *comscasis* from the Center for Ocean Mega-Science, Chinese Academy of Sciences).

Cells are Gram-stain-negative curve-shaped, 2.0-6.0 μm in length and 0.5-0.8 μm in width. Facultative anaerobic and oxidase-positive. The temperature range for growth is 28-37 °C with an optimum at 30 °C. Growing at pH values of 6.0-8.0 (optimum, pH 7.0). Growth occurs at NaCl concentrations between 0.0-5.0% with optimum growth at 1.0% NaCl. By analyzing the hydrolysis of polysaccharides, the growth is promoted significantly by cellulose, pectin and xylan. From the sole carbon source utilization test, growth is stimulated by acetate, maltose, fructose, lactate, sorbitol and D-mannose. Weak growth occurs with ethanol and formate. The major polar lipids are phosphatidylethanolamine, one unidentified phospholipid, one unidentified aminolipid, one unidentified lipid. Containing significant proportions (>10%) of the cellular fatty acids iso-C_15:0_, C_16:0_, summed feature 3 (containing C_16:1_ω7*c* and/or C_16:1_ω6*c*) and summed feature 8 (containing C_18:1_ω7*c* and/or C_18:1_ω6*c*).

The type strain, WC007^T^ (=KCTC 25169^T^ =MCCC 1K04777^T^), was isolated from the sediment of deep-sea cold seep, P.R. China. The DNA G+C content of the type strain is 38.38%.

