## Supplemental Figures and Tables for "*Maribellus comscasis* sp. nov., isolated from the deep-sea cold seep"

**Category:** New taxon in Bacteroidetes

**Running title:** A novel Bacteroidetes bacterium

The NCBI GenBank accession numbers for the 16S rRNA gene sequence and whole-genome sequence (WGS) of strain WC007^T^ are MN096653 and CP046401, respectively.

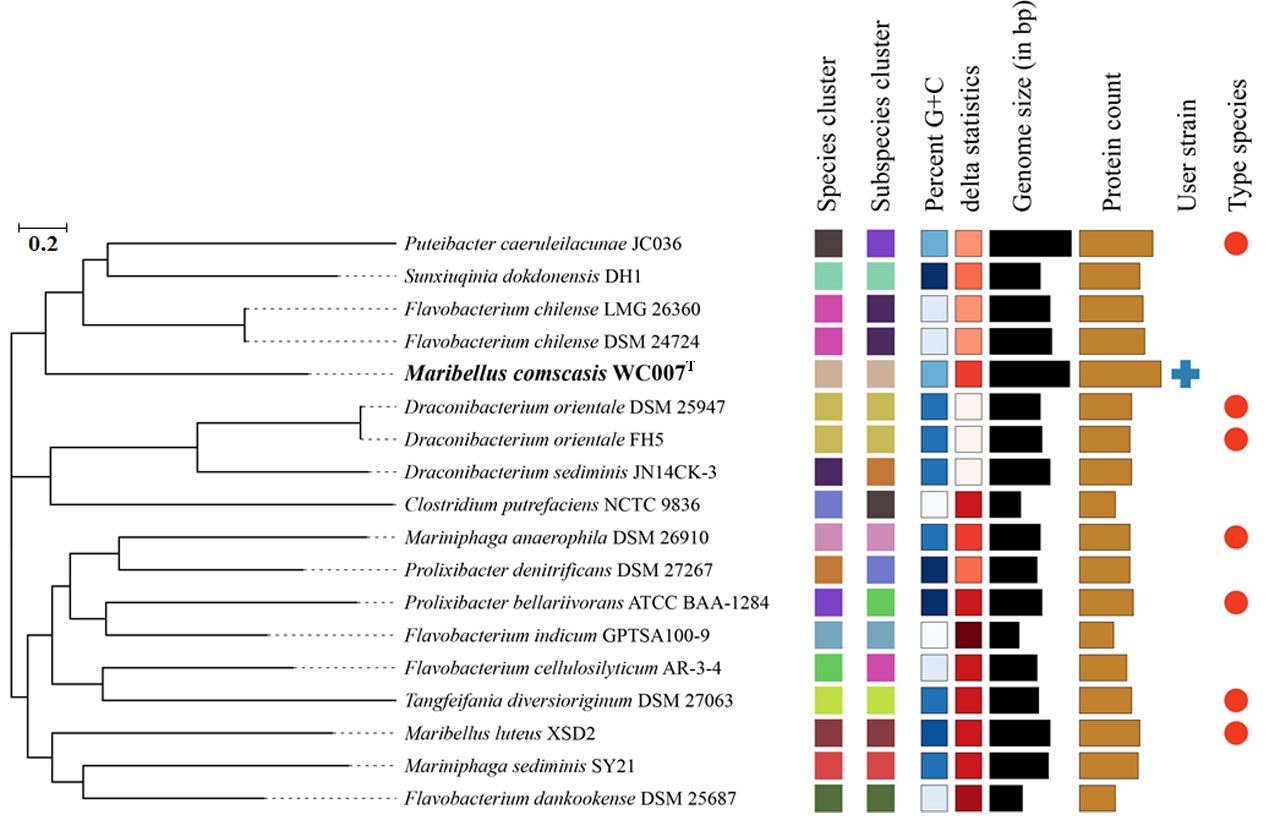
**Supplemental Figure 1.** Phylogenomic tree analysis of *M. comscasis* WC007^T^ based on the TYGS algorithm (https://tygs.dsmz.de/).

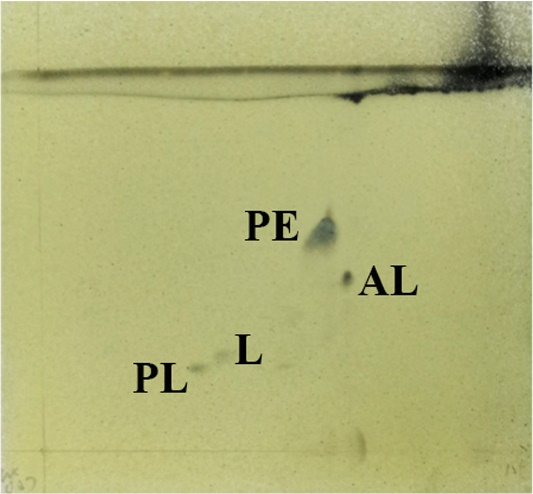

**Supplementary Figure 2. The polar lipids of strain WC007^T^ as revealed by two-dimensional TLC.** Chloroform/methanol/water (65:25:4, v/v/v) was used in the first direction, followed by chloroform/methanol/acetic acid/water (80:12:15:4, v/v/v/v) in the second direction. The plate was sprayed with 10% ethanolic molybdophosphoric acid. Abbreviations: PE, phosphatidylethanolamine; PL, one unidentified phospholipid; AL, one unidentified aminolipid; L, one unidentified lipid.

**Supplementary Table 1.** Genomic characteristics of strain WC007^T^ and other closely related strains.

| **Characteristics** | **WC007^T^** | **XSD2^T^** | | **SY21^T^** | | **FH5^T^** |
| --- | --- | --- | --- | --- | --- | --- |
| Gene Bank ID | CP046401 | QWGR00000000 | QWET00000000 | | CP007451 | |
| Genome size (bp) | 7,811,310 | 6,145,806 | | 5,826,160 | | 5,073,141 |
| No.scaffolds/contigs | 1 | 354 | | 70 | | 89 |
| GC-content (%) | 38.4 | 44.1 | | 41.7 | | 41.3 |
| ANIb (%) | 100 | 70.11 | | 70.28 | | 70.51 |
| ANIm (%)  AAI (%) | 100  100 | 84.94  71.0 | | 83.71  71.7 | | 84.35  71.3 |
| Tetra | 1 | 0.92022 | | 0.9496 | | 0.9397 |
| *is*DDH (%) | 100 | 20.40 | | 18.40 | | 20.20 |
